# Dissecting the Thermodynamics of ATP Binding to GroEL One Nucleotide at a Time

**DOI:** 10.1101/2022.09.13.507831

**Authors:** Thomas Walker, He Mirabel Sun, Tiffany Gunnels, Vicki Wysocki, Arthur Laganowsky, Hays Rye, David Russell

**Affiliations:** Department of Chemistry, Texas A&M University, College Station, TX 77843; Department of Chemistry and Biochemistry, The Ohio State University, Columbus, OH 43210; Department of Biochemistry & Biophysics, Texas A&M University, College Station, TX 77843

## Abstract

Understanding how large, oligomeric protein complexes respond to the binding of small ligands and other proteins is essential for describing the molecular basis of life. This in turn requires a complete characterization of the binding energetics and correlation of thermodynamic data with interacting structures, including effects of small molecules and solvent. However, the size of many protein oligomers, the myriad intermediate ligation states they can populate, and their often complex allosteric regulation typically restrict analysis by traditional methods to low resolution, ensemble averages. Here, we employ variable-temperature electrospray ionization native mass spectrometry to determine the thermodynamics for stepwise binding of up to ATP molecules to the 801 kDa GroEL complex, a tetradecamer chaperonin complex. Binding thermodynamics reveal strong enthalpy-entropy compensation (EEC) and high degrees of cooperativity are observed for formation of GroEL-ATP_7_ and GroEL-ATP_14_. These are evidenced by entropically favored ATP binding to the *cis* ring (formation of GroEL-ATP_1-7_), with variations in EEC for subsequent binding of ATP to the *trans* ring (GroEL-ATP_8-14_), as expected for negative inter-ring cooperativity. Entropy driven ATP binding to the GroEL tetradecamer is consistent with ligand induced conformational changes of the GroEL tetradecamer, though the magnitude of the entropy change suggests that reorganization of GroEL-hydrating water molecules and/or expulsion of water from the GroEL cavity may also play a key role. By determining the thermodynamic signatures for individual ligand binding reactions to the large, nearly MDa GroEL complex, we expand our fundamental view of chaperonin functional chemistry. Moreover, this work and related studies of protein-ligand interactions illustrate unparalleled capabilities of vT-ESI-nMS for thermodynamics studies of protein interactions with ligands, and other molecules, such as proteins and drugs.

## Introduction

Electrospray ionization native mass spectrometry (ESI-nMS) has evolved as powerful method for studies of protein-cofactor, protein-ligand, and non-covalent protein-protein interactions. MS-based strategies are attractive because they require only very small amounts of sample, permit product stoichiometries to be directly obtained, and allow reaction kinetics and thermodynamics can be quantified. Klassen and coworkers recently reported a new strategy, “quantifying biomolecular interactions using slow mixing mode (SLOMO) novel nanoflow ESI-MS”, for determination of equilibrium binding affinities (K_D_ at 25 °C) for biomolecular interactions,^1^ and they demonstrated the utility of this approach for a number of peptide- and protein-ligand systems. Interactions of proteins with cofactors, ligands, and other proteins often result in conformational changes and/or reorganization of solvent that result in substantial shifts in enthalpy (ΔH) and entropy (-TΔS). At the same time, solution temperature, pressure, and concentration can influence changes in the “native” structure of biomolecules through manipulation of these thermodynamic contributions.^2, 3^ Changes in thermodynamic contributions are often hallmarks of changing conformational states and vice-versa.^3^ Thermodynamic analysis is thus a powerful approach for probing fundamental mechanisms as well as for extracting critical information on structure/activity relationships that are important for drug discovery and drug design, including water-mediated interactions.^4–6^

Electrospray ionization produces ions by formation of nanodroplets from which solvent rapidly evaporates, cooling the nanodroplet contents to temperatures of 130-150 K.^7^ Beauchamp and coworkers described this evaporative drying process as “freeze-drying”.^7^ We have used this approach to track the structural evolution of hydrated biomolecules in route to forming solvent-free, gas-phase ions,^8^ and we, and others, have shown that this approach can be used to capture native and non-native protein states that co-exist in solution. There exist important parallels between cryo-EM ESI-MS/cryo-IM-MS (*vide infra*) in that both techniques take advantage of kinetic-trapping of molecules as they exist in solution.^9^

While traditional solution-based techniques can be used to robustly examine temperature-dependent interactions between biomolecules and their ligands, they generally report on the ensemble average of ligand bound states present in solution. Recent work leveraging the molecular resolution of nMS has shown that species-resolved thermodynamic analysis is possible.^10–12^ Combined with nMS, vT-ESI allows for thermodynamic measurements of solution-phase structures with the benefit of mass separation.^11, 13, 14^ This type of species-specific thermodynamic analysis can be especially valuable for complex or heterogeneous protein-ligand systems where the binding mechanism fundamentally changes as a result of a perturbation or shift in conditions without a measurable alteration to the observed Gibbs free energy (ΔG). In these cases, enthalpic and entropic contributions to the Gibbs free energy shift in opposite directions, a phenomenon known as enthalpy-entropy compensation (EEC). ^15, 16^

We previously reported EEC results for protein complex-lipid binding that varied with lipid head group, tail length and showed that mutant forms of AmtB altered phosphatidylglycerol (PG) binding site having distinct changes in the thermodynamic signatures.^10^ VT-nESI studies of lipid binding to the human G-protein-gated inward rectifier potassium channel, Kir3.2, displays distinct thermodynamic strategies to engage phosphatidylinostitol (PI) and phosphorylated forms thereof. The addition of a 4’-phosphate to PI results in an increase in favorable entropy. PI with two or more phosphates exhibits more complex binding, where lipids appear to bind to nonidentical sides on Kir3.2; interactions of 4,5-bibis-phosphate with Kir3.2 is solely driven by large, favorable entropy whereas adding a 3’phosphate to PI(4,5)P_2_ displays altered thermodynamics.^11^ The lipid acyl chain has a marked impact on binding thermodynamics, and in some cases, results in favorable enthalpy. More recent studies using vT-ESI nMS combined with ion mobility showed that the cysteine desulfurize enzyme IscU exists in structured, intermediate, and disordered forms that rearrange to more extended conformations at higher temperatures. Comparisons of Zn-IscU and apo-IscU reveals that Zn(II) binding attenuates the cold/heat denaturation of IscU, promotes refolding of IscU, favors the structured and intermediate conformations, and inhibits formation of the disordered high-charge states.^17^ Collectively, these studies highlight how vT-ESI-nMS can be applied as a powerful approach for studies of relationships among temperature, conformation and ligand interactions in a complex biomolecular system.

Molecular chaperones represent another large class of essential biomolecules for which ligand binding and conformational changes are intimately linked to function. The chaperonin family of molecular chaperones, or Hsp60s, are large oligomeric protein complexes that utilize the energy of ATP hydrolysis to actively facilitate protein folding. The canonical bacterial chaperonin, GroEL, is an 801 kDa tetradecamer protein complex from *E. coli* that consists of two heptameric stacked rings. Each GroEL subunit consists of three domains—apical, intermediate, and equatorial.^18–20^ The apical domain is highly dynamic and is responsible for binding protein substrates and the co-chaperonin GroES. The intermediate domain acts as a hinge between the equatorial and the apical regions of each subunit, and the equatorial domain of each subunit is the least dynamic and serves as the interfacial contact between each heptameric ring. ^21, 22^ The equatorial domain also harbors the ATP binding site for each subunit.^19^ While the structure, dynamics, and ATP binding of GroEL has been extensively investigated,^23, 24, 25–30^ a number of fundamental questions about how ATP binding and hydrolysis controls the GroEL functional cycle remain unresolved.

A particular limitation for addressing many of the fundamental issues is the complexity of the GroEL-ATP system, *i.e*. the tetradecameric complex can bind up to 14 nucleotides. At the same time, ATP binding to the GroEL tetradecamer is known to induce structural changes in the tertiary structure of the GroEL subunits while simultaneously rearranging the quaternary structure of the oligomer.^25, 26^ It has been shown via X-ray crystallography and cryo-EM that the binding of ATP by GroEL induces an extension and twisting of the apical domain^30^ and a small “rocking” of the equatorial domain.^6, 26^ The binding of ATP by GroEL is also influenced by the presence of cations (*e.g*., Mg^2+^ and K^+^). Mg^2+^ is necessary for the binding of ATP and K^+^ is thought to activate the ATPase activity of GroEL.^31^ It has also been proposed that NH_4_^+^ ions can act as a surrogate for K^+^ ions.^32^ These nucleotide-driven structural rearrangements interact to create a complex and layered set of allosteric transitions that govern the GroEL protein folding cycle. This ligand binding and structural complexity constrains the detail that can be robustly extracted from ensemble thermodynamic studies.

Recently, we reported results using native mass spectrometry (nMS) that reveal new insights about GroEL oligomer stability and the stoichiometry of GroEL-ATP/GroES interactions.^33^ Using variable-temperature electrospray ionization (vT-ESI) we found that GroEL-ATP binding was temperature and ATP-concentration dependent, suggesting that more detailed thermodynamic analysis might reveal new insights into how the GroEL nanomachine operates. Here we report the thermodynamic measurements (ΔH, ΔS, and ΔG) of ATP and ADP binding to GroEL utilizing nMS with vT-ESI, along with studies that observe how endogenous ions affect the GroEL-ATP binding interaction. We demonstrate that this approach can extract rich thermodynamic measurements for stepwise binding of ATP to GroEL. We further show that EEC plays a central role in the binding of ATP to a GroEL ring, with important mechanistic implications for how GroEL functions. These results thus lay the groundwork for the development of new strategies for detailed thermodynamic analysis of other complex, multi-dentate biomolecular systems. The ability to measure and understand the underlying thermodynamics of protein-cofactor, protein-ligand and proteinprotein interactions has been the focus of much research particularly in drug discovery efforts.^34^

## Results and Discussion

### Thermodynamics of GroEL-ATP binding in ethylenediammonium diacetate (EDDA) buffer

**Figure 1A** contains deconvoluted mass spectra obtained for solutions of GroEL containing magnesium acetate and EDDA taken at temperatures of 5 °C, 23 °C, and 41 °C, and **Figure 1B** shows the intrinsic equilibrium constants (K_a_) for each GroEL-ATP binding reaction. The intrinsic binding constants are statistically corrected to account for the number of modes in which ligands may associate or dissociate from the complex based on the distribution of available monomers (see methods).^35^ It is interesting to note that the binding of up to GroEL-ATP_14_ is only observed at lower temperatures for 25 μM ATP and that binding affinity is decreased at higher temperatures. The deconvoluted mass spectra also reveal potential cooperativity in binding at lower temperatures, *e.g*., note the increase in the abundance of GroEL-ATP_14_ at 5 °C. The association constants represent the sequential binding of ATP to GroEL. The K_a_ values vary significantly with solution temperature with binding of ATP being favored for each reaction at 5 °C compared to 41 °C, except for GroEL-ATP_13_. Most pronounced of all the constants, however, is GroEL-ATP_14_ where the temperature dependance is most easily observed. A similar effect for the GroEL-ATP_14_ K_a_ value was observed by Sharon *et al*., albeit their study was only conducted at room temperature.^36^ A depression in the GroEL-ATP_8-11_ K_a_ values are hallmark of negative inter-ring cooperativity and is most apparent at low solution temperatures.

**Figure 1.**
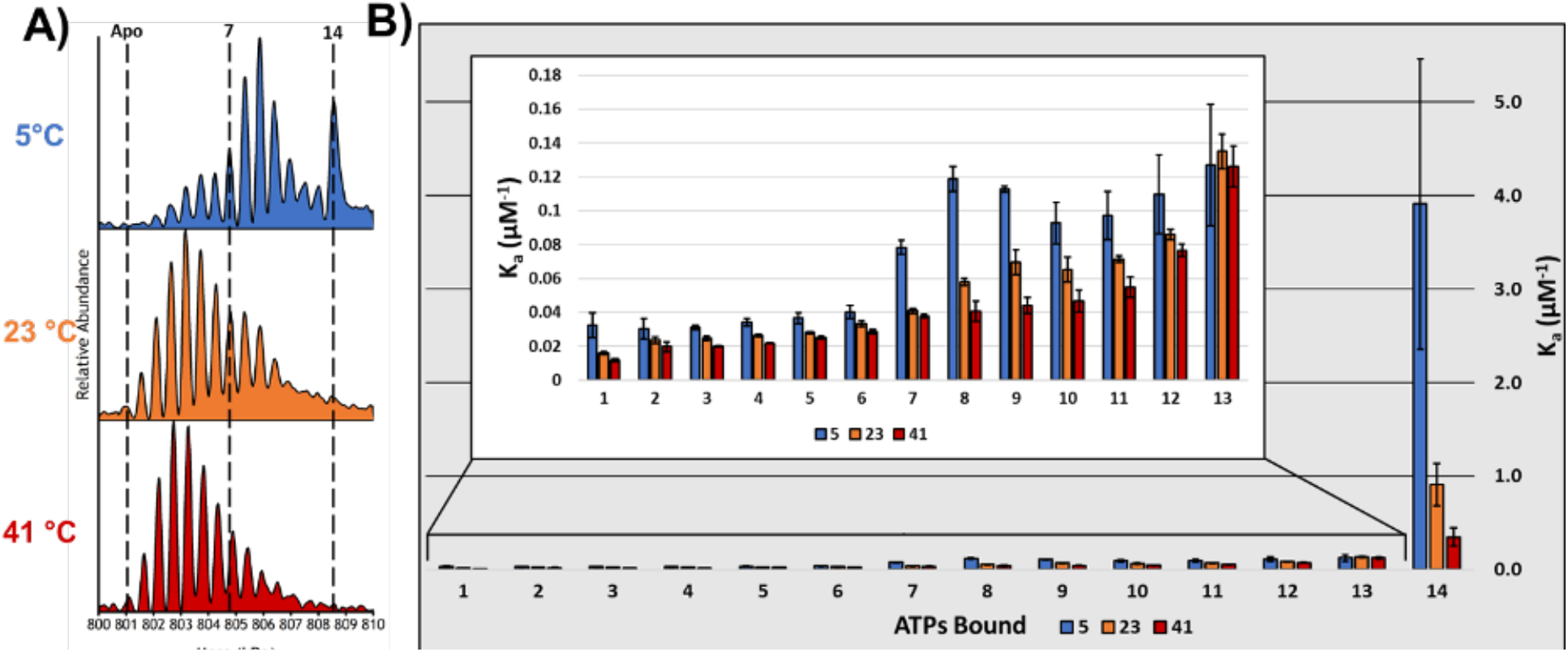
**A)** Deconvoluted mass spectra of 500 nM GroEL in a solution of 200 mM EDDA, 1 mM MgAc_2_, and 25 μM ATP at 3 different temperatures. Binding affinity diminishes as solution temperature is increased. Bimodality in the ATP binding distributions is likely to be a consequence of negative cooperativity between the *cis* and *trans* rings. **B)** This bar charts shows the intrinsic K_a_ values calculated for the 14 ATP binding reactions for GroEL at 3 different temperatures 5 °C, 23 °C, and 41 °C. The inset shows a bar chart that expands the first 13 binding reactions so that details may be more observed easily. Binding affinity is temperature dependent and colder solution temperatures enhance the effect of inter-ring negative cooperativity, as the affinity for ATP decreases more substantially when binding in the *trans* ring begins.

**Figure 2A** contains deconvoluted MS spectra for concentration-dependent ATP binding at 25 °C for 10, 25, and 50 μM ATP in EDDA and the absence of magnesium, which prevents GroEL from turning over ATP. These results are consistent with those reported by Cliff *et al.;* low concentrations of ATP promote binding to the cis ring to form GroEL-ATP_7_, whereas higher concentrations promote ATP binding to both rings to form GroEL-ATP_14_. ^37^ The bimodal binding patterns for observed at 23 °C (**Figure 1A**) and also for solutions containing 25 μM ATP (**Figure 2A**) may be a result of rearrangement reactions they attributed to structural transitions preceding hydrolysis. Van’t Hoff analysis of the data shown in **Figure 1B** was used to evaluate the thermodynamics (ΔG, ΔH and TΔS at 25 °C) for each ATP binding reaction that are plotted in **Figure 2B** and **2C** (see SI for details). The ratios for ΔH and -TΔS illustrate a high degree of EEC compensation, which in most cases reflect entropy driven ATP binding. At low temperature (5 °C) the ΔG values show the most diverse pattern for binding of ATP potentially displaying inter-ting negative cooperativity (**Figure S1**). ΔG values in **Figure 2B** for *cis* ring ATP binding range from −24.0 to −26.4 kJ·mol^-1^ and becomes more favored (<-27.1 kJ·mol^-1^) for GroEL-ATP_8-13_. The ΔG terms become more favorable with each additional ATP bound before becoming disproportionately favored (−33.7 kJ·mol^-1^) for GroEL-ATP_14_. As the solution temperature increases, these variations in ΔG diminish. ΔΔG values for 5 - 41 °C (**Figure S2**) reveal that binding to the *cis* ring (GroEL-ATP_1-7_) becomes more favored as the temperature is increased when compared to the initial (GroEL-ATP_8-10_) ATP binding reactions of the *trans* ring.

**Figure 2.**
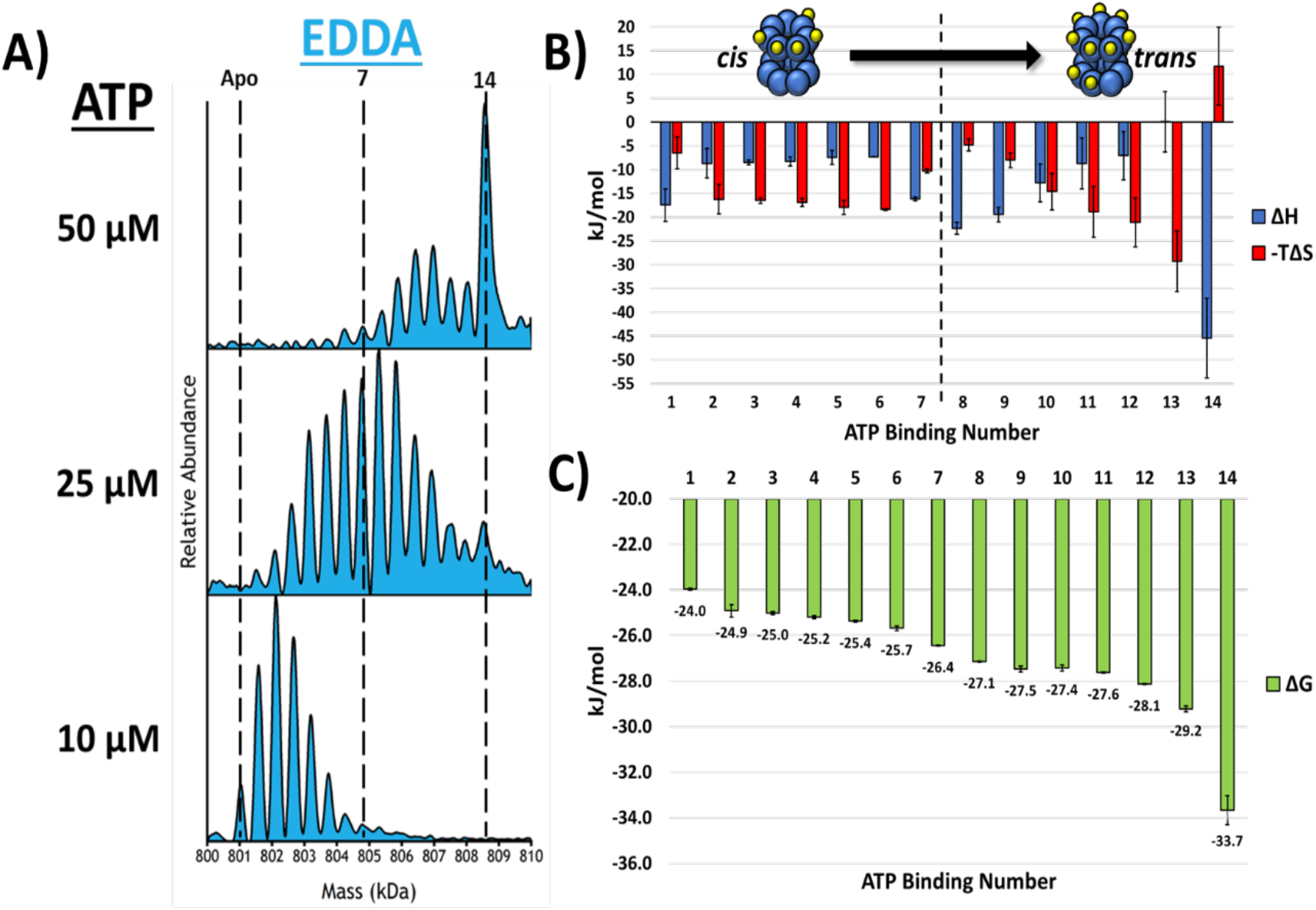
**A)** Deconvoluted spectra showing the effects of increase in ATP concentration. **B)** The histograms show enthalpy (ΔH) and entropy (-TΔS) for the individual ATP binding reactions at 25 °C. The observed EEC for the first 7 binding reactions is quite different from that for binding of 8-14 ATPs, which is consistent with filling of *cis* ring prior to binding to *trans* ring. Note also that EEC is drastically different for addition of the 13^th^ and 14^th^ ATP cofactor. These observed changes in EEC are indicative of substantial structural changes in the of GroEL complex. **C)** The overall changes in ΔG associated with each of the 14 ATP binding reactions. All error bars are the standard deviation of 3 replicates.

The changes in enthalpy and entropy (EEC) are strong evidences for conformational changes associated with binding of ATP to GroEL.^15^ These changes in EEC are a phenomenon that as ΔH or TΔS varies the other tends to compensate in an opposite direction (see **Figures 2C**, **3C**, and **S3**); this effect has been observed in thermodynamic measurements of other interactions. The entropy term encompasses both the conformational entropy of the GroEL complex but also the entropy of the solvent,^38^ and the enthalpy term is subject to the same contributions of structure and solvent.^39^ For example, in the GroEL system binding of ATP leads to extension of the apical domain and release of confined water, both of which are entropically favorable.^4–6^ However, a more extended, labile apical domain obviates a loss of favorable binding contacts (Van der Waals, H-bonding, and salt bridging) which would be enthalpically less favorable. The resultant ΔH value for that interaction would be increased and compensated by a decrease in -TΔS, *i.e*., become more favorable. Changes in the structure of GroEL complex are expected to alter the hydration of the complex as well as confined water in the GroEL cavity. Similar change in confined water has been reported by Brown *et al*. for rhodopsin.^40^ Csermely described similar effects of confined water in his “chaperone-percolator model,” specifically influx of water into the GroEL cavity and/or efflux of confined water, both of which are expected to be entropically favored and consistent the EEC trends shown in **Figure 2B**.^41^

**Figure 3.**
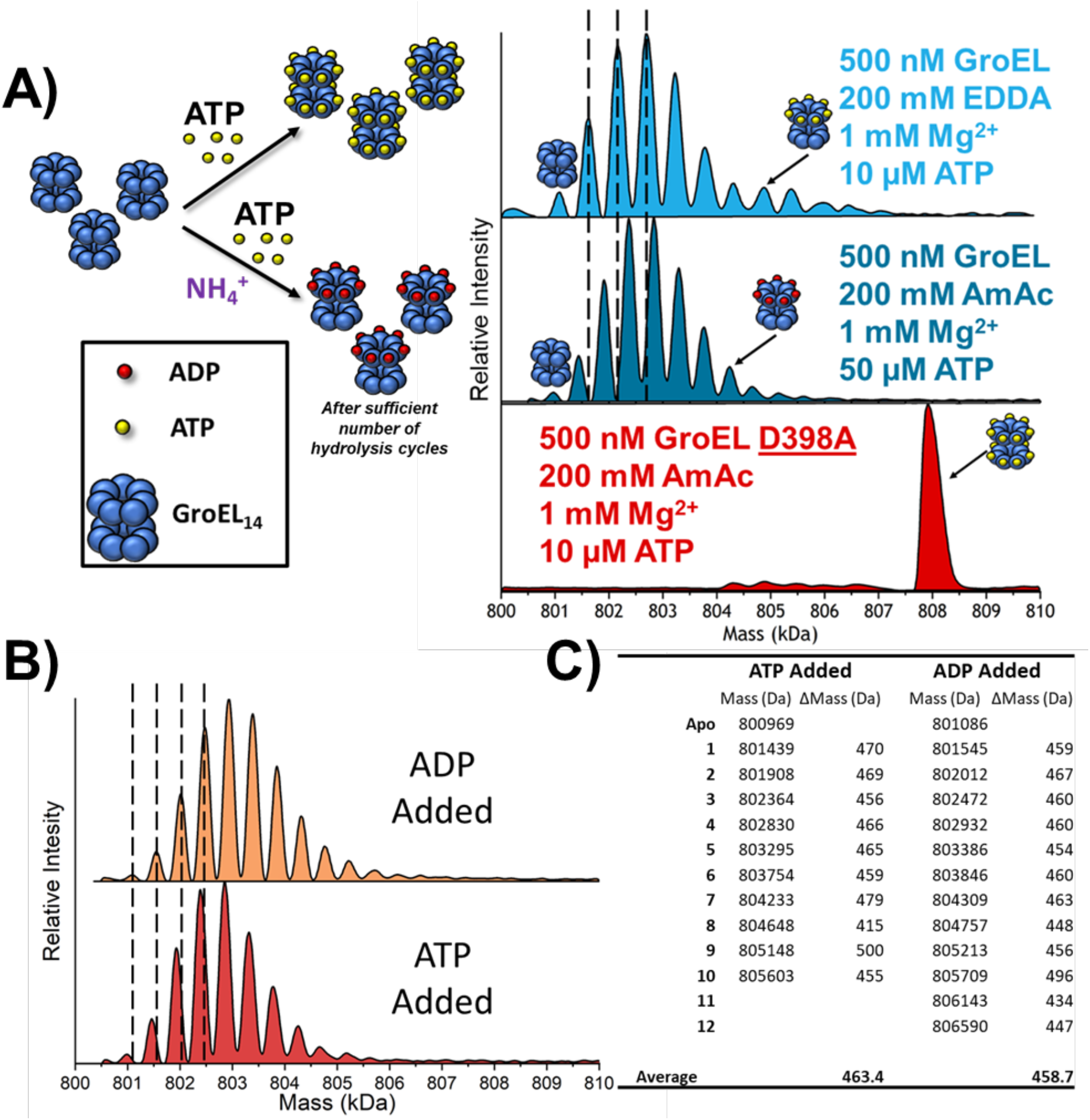
**A)** Stacked spectra showing the difference in mass shifts observed for nucleotide binding observed in EDDA and AmAc solutions. Black lines are used to aid the viewer and show that the mass shifts for EDDA are assigned to [ATP + Mg^2+^]_n_ while the mass shifts for AmAc conditions are assigned to [ADP + Mg^2+^]_n_. A hydrolysis deficient mutant GroEL^D398A^ was also analyzed in an AmAc solution (red) which shows the elevated level of cooperative binding in the presence of NH_4_^+^ ions and the absence of hydrolysis. Also note for GroEL^D398A^ that the affinity and cooperativity of the GroEL mutant for ATP is drastically increased in AmAc compared to EDDA conditions. **B)** Stacked deconvoluted mass spectra showing the similarities in the binding distributions when either ADP is directly added to a solution containing GroEL, or ATP is added under conditions where hydrolysis occurs. Solution conditions are 500 nM GroEL, 50μM ATP or ADP, 1mM MgAc_2_, and 200 mM AmAc at 25 °C. **C)** Table containing the peak centroid data for **A**. Mass shift values are in the form Δ*Mass* = (*n* + 1) – *n*. **Note**: measured mass of apo complex for ATP vs ADP is shifted by about 120 Da explaining why subsequent peaks are not exactly aligned in panel **B**.

EEC observed for ATP binding explains the comparative lack of ΔG fluctuation across the range of ATP binding reactions. With the exception of the first and last ATP binding reactions, binding of ATP to the *cis* ring of GroEL is largely entropically driven and is indicative of structural rearrangement. The observed deviations for EEC in the region of GroEL-ATP_7-10_ coincides with the filling of the *cis* ring and the transition to binding in the *trans* ring. These values switch again for GroEL-ATP_11-13_ and yet again for GroEL-ATP_14_. The binding of the ATP_14_ is the only binding reaction with small unfavorable entropy. This seems to indicate that the final structural rearrangement in the *trans* ring is very ordered compared to preceding binding reactions (GroEL-ATP_11-13_) and is very favored enthalpically which explains its highly favorable ΔG value.

### Effects of NH_4_^+^ ions on GroEL-ATP binding

In previous work we showed that that temperature has strong effects on GroEL-ATP binding as well as the overall stability of the GroEL tetradecamer. Of particular note was the pronounced effects for AmAc compared with EDDA solutions, as well as differences in the GroEL tetradecamer average charge states, Z_avg_ of 66^+^ versus 56^+^, respectively. Lorimer and coworkers showed that K^+^ ions are necessary for the activation of the ATPase mechanism of GroEL.^42, 43^ Seidel^44^ and Lorimer also report evidence that NH_4_^+^ ions can act as K^+^ surrogates, which suggests that the observed effects of the ESI buffers may be linked to the initiation of ATPase activity in GroEL. This possibility was tested by addition of ATP to two solutions with one containing GroEL in AmAc and the other GroEL in EDDA. Deconvoluted mass spectra in **Figure 3A** show that mass shifts for nucleotide binding are about 80 Da smaller in AmAc than in EDDA (mass shifts of 460 Da correspond to [Mg^2+^ + ADP]). The observed mass shifts in AmAc solution are clear evidence for GroEL driven hydrolysis of ATP. When compared to experiments conducted in EDDA buffer, it is clear that NH_4_^+^ is necessary for GroEL ATPase activity (**Figure 3A**). We also compared the effects of AmAc on hydrolysis by collecting data for the hydrolysis deficient GroEL D398A mutant (GroEL^D398A^), which only [Mg^2+^ + ATP] was observed bound to the complex (**Figure 3A**). It is interesting to note that for GroEL^D398A^, the presence of NH_4_^+^ ions greatly increase the binding affinity and cooperativity of binding ATP. Lastly, we also compared solutions of GroEL in AmAc solutions that contained ADP or ATP and for both solutions the ions detected solely corresponded to ADP bound complexes (**Figure 3B** and **C**).

### Thermodynamics of GroEL-ATP/ADP binding in ammonium acetate (AmAc) buffer

Ammonium acetate is a commonly used native MS buffer, but monovalent ions (K^+^ and Rb^+^) are known to catalyze GroEL ATP hydrolysis.^43, 45^ The capability to measure individual ATP binding reactions coupled with vT-nESI thermodynamic measurements is an excellent way for determining whether monovalent ions are directly or indirectly linked to the effects of ATP binding and subsequent hydrolysis. In AmAc solutions we only observe GroEL-ADP products (*vide infra*). **Figure 4A** contains the deconvoluted mass spectra for ADP concentration-dependent binding to GroEL. When compared to **Figure 2A**, it is clear that GroEL binds fewer ADP than ATP (in EDDA), moreover cooperative binding of ADP is not apparent. **Figures 4B** and **4C** show thermodynamic data for GroEL-ADP*n* binding the *n^th^* ADP molecule at 25 °C; the thermodynamic constants were only calculated up to GroEL-ADP_9_ due to poor fits for the Van’t Hoff plots for GroEL-ADP_10-14_. Note the changes in EEC for GroEL-ADP_4-5_ compared to that for GroEL-ADP_1-3_. The comparison of Gibbs free energy values between ADP and ATP binding also demonstrates that ADP binds more weakly than ATP in EDDA (see also **Figure S4**). In EDDA the ΔG for the binding of ATP at 25 °C is about 2-3 kJ·mol^-1^ more favorable than for ADP binding in AmAc conditions.

**Figure 4.**
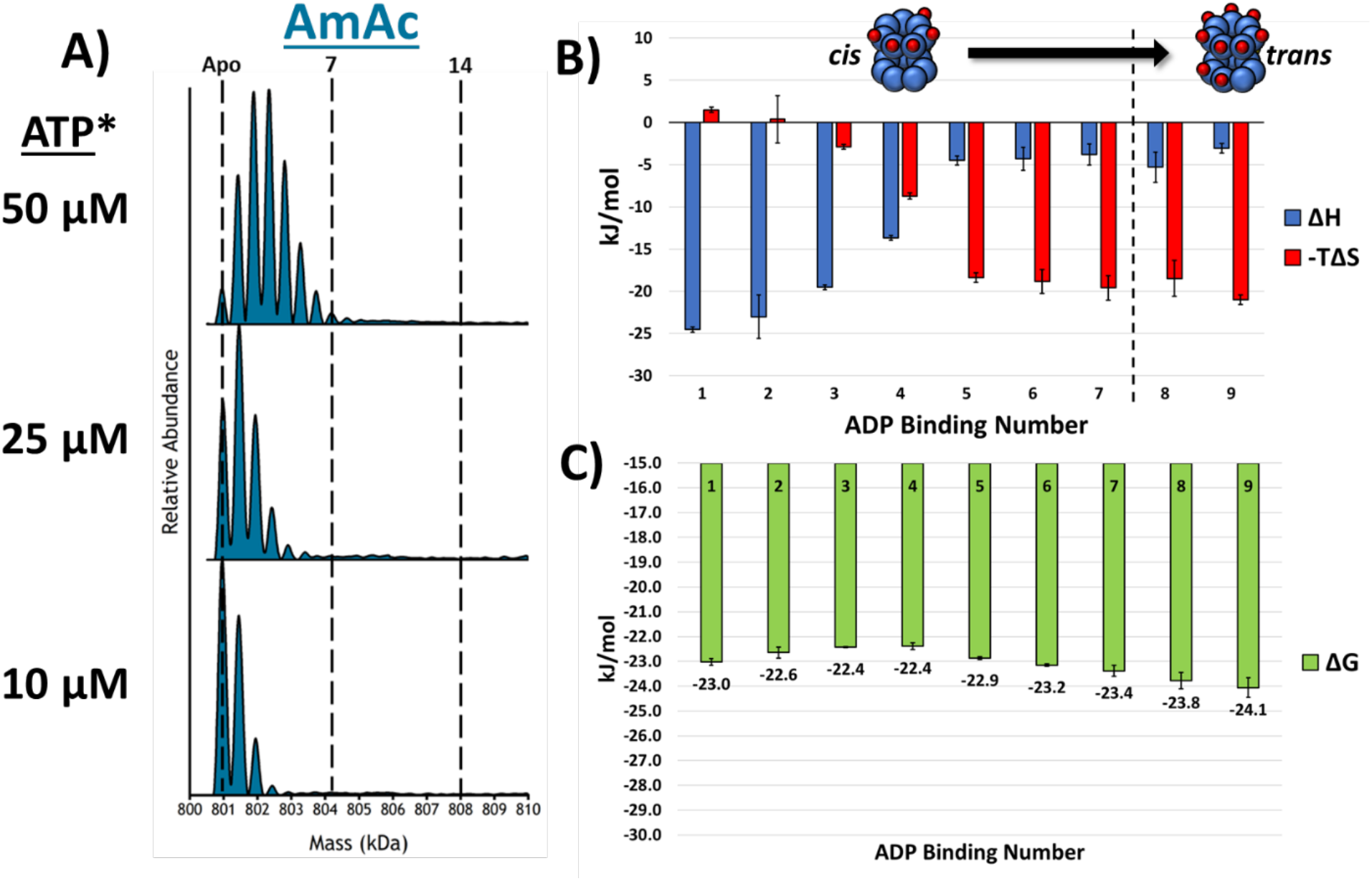
**A)** These stacked deconvoluted spectra show that as ADP concentration is increased the binding of ADP is does not show any cooperativity. **B)** Bar chart showing the ΔH and -TΔS contributions at 25 °C for each of the ADP binding reactions. The enthalpy and entropy show a singular transition from GroEL-ADP_4-5_and are overall much less dynamic than the binding of ATP in EDDA. **C)** This bar chart displays the Gibbs free energy measurements for the ADP binding reactions at 25 °C. EEC is more heavily present in the ADP data set as the Gibbs free energy varies by less than 2 kJ/mol. All error bars are the standard deviation of at least 3 replicates. *(**Note:** ATP was added to the solution but only ADP binding was observed.)

As shown in **Figure 1**, the ATP binding affinity of GroEL in EDDA is temperature dependent, and changes in the ΔG values for the *cis* ring binding reactions is of greater magnitude than for several of those in the *trans* ring (**Figure S2**). The initial ATP binding to the *cis* ring is more favorable compared to the *trans* ring binding reactions, which are more allosterically disfavored as evidenced by subtle differences in symmetry between the rings observed in ATP-bound cryo-EM structures.^23^ A potential structural explanation concerns the inter-ring hydrophobic contacts in the region of A109 on helix D that are extensively perturbed by the binding of ATP in the binding pocket of the equatorial domain.^20, 29^ The disturbance of these contacts is theorized to be responsible for inter-ring negative cooperativity upon the binding of ATP.^6, 29, 46^ At higher temperatures increased dynamics in this region would only serve to further weaken this contact, and potentially increase the prevalence of the negatively cooperative mechanism.

The data in **Figure 1A** show that as solution temperature is increased, the binding affinity for ATP diminishes. It is interesting to note the behavior of the profile for the deconvoluted spectra; at colder solution temperatures, there exists a bimodal distribution of ATP bound to the complex. The two distributions coincide with the filling of each heptameric ring of GroEL. Horovitz *et al*. demonstrated that the binding of ATP to GroEL can be explained through a nested-cooperativity model where each ring binds concertedly (Monod-Wyman-Changeux (MWC)) and the behavior between the rings is coupled sequentially (Koshland-Nemethy-Filmer (KNF)).^47^ Binding of GroEL-ATP_14_ observed in the MS data can be interpreted as a highly cooperative reaction as the relative intensity of the GroEL-ATP_14_ signal is largely disproportionate compared to the preceding distribution of signals (GroEL-ATP_8-13_). As the *trans* ring begins to bind several ATP molecules (GroEL-ATP_8-10_) the cooperative effect allosterically begins to favor binding of a full ring. At higher temperatures much of this cooperative signature is lost as binding of ATP to the *cis* ring becomes favored.

The data contained in **Figure 2B** provides clear evidence that formation of GroEL-ATP_1-7_ (EDDA) is largely driven by entropy, which is consistent with the known rotation of the “hinge” region of the intermediate domain and extension of the apical domain initiated by loss of salt bridge contacts (E255-K207 and R197-E386) that connect adjacent apical domains in the subunits of the apo-GroEL.^6, 26^ The enthalpy and entropy terms for binding GroEL-ATP_7-10_ switch as the binding becomes mostly enthalpically driven. These differences in EEC are expected owing to negative cooperativity associated with inter-ring communication accompanying GroEL-ATP binding. Saibil and coworkers report that upon ATP binding to the *cis* ring of GroEL the equatorial domain shifts resulting in loss of a key hydrophobic contacts.^6, 29^ The loss of these contacts is thought to be the structural progenitor of the inter-ring negative cooperativity and may explain why entropy and the overall favorable Gibbs free energy is decreased for the initial ATP binding to the *trans* ring (**Figures 2B**, **S1**, **S2**). Entropy then dominates for the formation of GroEL-ATP_11-13_, which is most likely associated with similar structural changes observed for ATP binding in the *cis* ring. This explanation is consistent with observations made by Chapman *et al*., viz. increased binding affinity when up to 3 ATP molecules are bound to a ring.^27^ The GroEL-ATP_14_ binding reaction coincides with the most favorable binding and is the only binding reaction that has an associated negative, unfavorable entropy. The GroEL-ATP_14_ binding reaction is largely enthalpically favorable because the restructuring of the complex occurs during the formation of GroEL-ATP_10-13_ making GroEL-ATP_14_ more enthalpically favored at the cost of an entropic penalty.

Another interesting point that has arisen during our thermodynamic studies is the clear differences observed in the thermodynamic signatures for the *cis* and *trans* rings of GroEL. The *cis* ring ATP binding is entropically favored whereas the *trans* ring ATP binding is much more diverse in its thermodynamic contributions. The binding of GroEL-ATP_1-7_ is entropy favored owing to the large conformational change in the apical domains, while the initial ATP binding reactions (GroEL-ATP_8-10_) in the *trans* ring are disfavored entropically; however, following binding of up to 3 ATP molecules to the *trans* ring the entropy and enthalpy switch for the subsequent binding reactions up to GroEL-ATP_13_.

The concerted (MWC) model of ligand binding postulates that the binding of one ligand to a protomer (subunit) induces changes in affinity for all the other protomers collectively into a relaxed or tense state.^48^ In contrast, the sequential (KNF) model argues that the binding of a ligand to a subunit will only have immediate effect (positive or negative) on its neighboring subunits.^49^ The thermodynamic signature for the GroEL-ATP_1_ binding reaction shows that it is enthalpically driven but the GroEL-ATP_2-6_ binding reactions are driven entropically and have entropic values that are relatively unchanging. This thermodynamic pattern in conjunction with the mass spectral data suggest that binding in the *cis* ring is largely sequential in the EDDA solution. The GroEL-ATP_1_ reaction sets into effect a “chain reaction” that affects adjacent subunits via allosteric transitions. If the binding were purely concerted, then the binding distribution should be like that seen in the *trans* ring where once 2 or 3 ATP molecules are bound then the distributions heavily shift to the filled state (GroEL-ATP_14_). These observations may be due to the difference in solution conditions used when comparing to solution-phase experiments. In these thermodynamic experiments there is no added K^+^ or GroES, which will ultimately change the affinity for ATP and conformation of GroEL. However, the observations made by the authors do support similar effects reported by previous investigators; specifically, the observation of cooperative binding of ATP that is non-stochastic.^5, 21, 47, 50^ Therefore, it is possible that the binding of ATP to GroEL is still governed via a nested-cooperativity model, but that the *cis* ring binding is governed sequentially, *trans* ring binding is concerted, and inter-ring communication is sequential but is negatively cooperative.

Comparison of the thermodynamic data for the binding of ADP in AmAc with the data for binding of ATP in EDDA reveals distinct differences between the two nucleotides. The presence of EEC at 25 °C causes the ΔG values in **Figures 2C** and **4C** to remain relatively constant. However, the underlying ΔH and TΔS values show a stark mechanistic difference between the two interactions. Initial binding reactions (GroEL-ADP_1-4_) for ADP-binding are driven enthalpically and not until GroEL-ADP_5_ does entropy become dominant. In the EDDA/ATP data, entropy becomes dominant much sooner (GroEL-ADP_2_) and several switching transitions occur as a function of sequential ATP binding. The lack of various switching reactions in the AmAc/ADP data confers an overall lack of cooperative binding of ADP; this conclusion is strengthened by the nMS data showing gaussian distribution for ADP binding at all observed ADP concentrations and by studies conducted by other researchers observing lower affinities associated with the binding of ADP.^28, 51^ It has been shown that binding of ADP can cause conformational shifts in GroEL,^52^ which are corroborated possibly by the entropic domination seen for GroEL-ADP5-9. However, these entropic shifts do not seem to be resultant from an allosteric transition^37^ that leads to increased affinity for further ligation reactions.

## Conclusion

Variable-temperature (vT) ESI native mass spectrometry (nMS) is a relatively new approach that complements isothermal titration calorimetry for studies of the effects of solution temperature protein-ligand interactions (ΔG, ΔH and TΔS). Furthermore, the vT-ES-nMS approach affords capabilities for determination of these quantities for individual binding reactions. Here, we demonstrate the utility of vT-ESI-nMS through determination of the thermodynamics for sequential ATP binding to the GroEL tetradecamer. By measuring these reactions as a function of temperature we are able to determine equilibrium association constants (K_a_) followed by Van’t Hoff analysis for determinations of ΔG, ΔH and - TΔS for each of the (ATP)_1-14_ binding reactions. The observed differences for *cis* and *trans* ATP binding exemplify nested cooperativity in which intra-ring binding is concerted (MWC) and inter-ring communication is sequential (KNF).^53^ These mechanistic differences are also revealed by differences in entropy, *cis* ring ATP binding is highly entropically favored whereas EEC for trans ring binding is highly variable. It is especially interesting to note the difference in EEC for ATP binding in EDDA versus AmAc buffers, cis ring ATP binding in AmAc solution largely enthalpy favored, but entropy favored for the trans ring. In both cases, the free energy changes for binding in a given ring are quite small, but variable between the two rings. Difference for enthalpy and entropy associated with binding of the last (14^th^) ATP in EDDA and (13 and 14th) in AmAc should not go unnoticed, highly enthalpically-driven in EDDA but entropically-driven in AmAc. The entropy components are indicative of structural changes of complexes and/or hydrating water network, whereas change in the enthalpy (and the overall free energy in the AmAc buffer) reflect changes in stability of the complex.

These thermodynamic data show complex patterns for enthalpy-entropy compensation (EEC) for GroEL-ATP binding. It is especially noteworthy that different thermodynamic mechanisms are observed for ATP binding to the *cis* and *trans* rings, *viz*. formation of GroEL-ATP_1-7_ and GroEL-ATP_8-14_, and the observed EEC also show significant dependences on the native ESI-MS buffers, ethylene diammonium diacetate (EDDA) and ammonium acetate (AmAc). In part these differences arise owing to hydrolysis of ATP to ADP in AmAc buffer, which is not observed in EDDA buffer; ATP hydrolysis in solutions containing NH_4_^+^ ions are thought to be analogous to that observed in solutions containing K^+^ ions. The MS data also support the existence of significant cooperative binding of ATP in EDDA solutions, which is enhanced further in AmAc solutions for hydrolysis deficient GroEL^D398A^. The synergistic effects of monovalent cations in binding of ATP and ATPase activity in GroEL remain an interesting aspect that necessitates further investigatin to elucidate their role in the manipulating the free energy landscape available to chaperonin complexes.

## Methods

### Sample Preparation

GroEL tetradecamer GroEL^D398A^ tetradecamer were expressed and purified by the Rye research lab at the Texas A&M Department of Biochemistry and Biophysics. Aliquots of the GroEL samples were stored at - 80 °C in a Tris buffer. Aliquots were buffer-exchanged into ammonium acetate (AmAc) or ethylenediamonium diacetate (EDDA) (obtained from Sigma-Aldrich) using BioRad biospin P-6 size exclusion (6000 Da cutoff) columns to remove unwanted salt contamination. Magnesium acetate (MgAc_2_) and Na-ATP were obtained from Sigma-Aldrich and fresh solutions were prepared prior to each experiment.

### Experimental

Data was collected on a Thermo Q Exactive UHMR (ultra-high mass range) mass spectrometer. Constituents for each sample were mixed immediately prior to analysis. For the thermodynamic analysis of GroEL-ATP binding solution conditions were: 1 mM MgAc_2_, 200 mM EDDA, 500 nM GroEL (14mer), and varying concentrations of ATP. The vT-nESI device was used to modulate the temperature of the solution; more information pertaining to operation of the device can be found in previous work.^13^ Solution temperatures used for this study were 5 °C to 41 °C; above 41 °C degradation products of the GroEL complex begin to become observable. The resolution setting was maintained at 25000, with 5 microscans, and injection time of 200 ms, capillary temperature was 150 °C, trap gas pressure was set to 7.0 (N_2_), desolvation voltage (in-source trapping)^54^ was set to -200 V, and HCD energy was set to 200 V (the latter two energy parameters only apply to EDDA buffer conditions). Care was taken to ensure that gas phase stability of the GroEL complex was retained; no monomer loss was observed for the energy setting listed previously. Also loss of nucleotide ligands is highly unlikely due to reports that nucleotide binding in the gas phase is irreversible.^55^ This was tested experimentally at high collision energies in which loss of ATP-bound monomers of GroEL were detected, signifying that the complex will dissociate prior to loss of nucleotide ligands in the gas phase (data not shown). Acquisition times for each spectrum were set to 1 minute. Under these conditions the ATP-bound states of GroEL were nearly baseline-resolved in most circumstances (**Figures S6** and **S8**). Thirteen solution temperatures, every 3 °C from 5 °C to 41 °C at 8 ATP concentrations (0, 1, 5, 10, 15, 25, 35, and 50 μM ATP) were analyzed in *n* = 3 trials. Each trial entailed the preparation of new solutions for buffers, GroEL, MgAc, and ATP solutions.

### Data Processing

Each spectrum was deconvoluted using UniDec^56^ and incorporated the 4 most abundant ATP distributions (see **Table S1** in supporting information for assignment statistics). Baseline reduction was avoided in the UniDec software as to not bias the data. The area of each peak to be integrated was determined by extrapolating the local minimum between peaks down to baseline to serve as the limit of integration. The resulting relative abundances were used in a sequential binding model (**Figure S5**) to fit K_d_ (dissociation constant) values.^10, 57^ The reciprocal of the K_d_ yields the K_a_; the natural logs of the K_a_ values were plotted against inverse temperature (in K) for Van’t Hoff analysis (**Figures S7** and **S9**). The slope of the fit line is used to calculate ΔH and the y-intercept is used to calculate ΔS in accordance with the equation 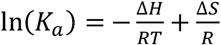. Using Δ*G* = Δ*H* – *T*Δ*S* the Gibbs free energy terms were calculated in units of 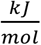.

## Supporting Information

More thermodynamics data along with peak assignment data, examples of raw spectra, and Van’t Hoff plots.

## Acknowledgment

Funding for this work was provided by the National Institute of Health grants P41GM128577 (D.H.R.; V.W.) and R01GM138863 (D.H.R.; A.L.)

## For Table of Contents Only

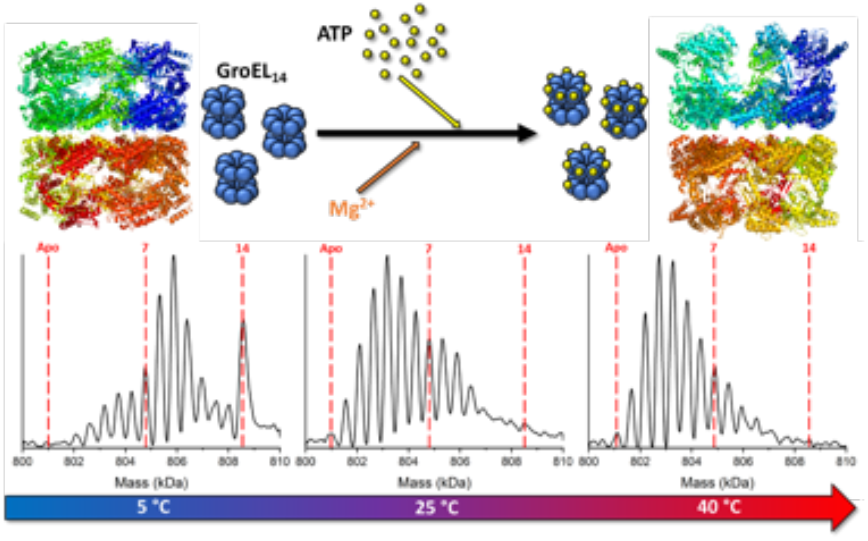

### Synopsis

A novel native mass spectrometry method for determining the thermodynamics of ligand binding to protein complexes, *viz*. enthalpy-entropy compensation for the binding of ATP and ADP to the chaperonin GroEL.

