## Supporting Information for "Dissecting the Thermodynamics of ATP Binding to GroEL One Nucleotide at a Time"

**Table of Contents**

| | | Page |
|---|---|---|
| **Figure S1** | **Change in ΔG in response to change in temperature for GroEL-ATP$_n$ in EDDA** | **S2** |
| **Figure S2** | **ΔΔG for GroEL-ATP$_n$ in EDDA and GroEL-ADP$_n$ in AmAc** | **S3** |
| **Figure S3** | **Comparison of EEC for ADP and ATP binding reactions** | **S4** |
| **Figure S4** | **Change in ΔG in response to change in temperature for GroEL-ADP$_n$ in AmAc** | **S5** |
| **Table S1** | **ATP assignment statistics** | **S6** |
| **Figure S5** | **Example plots used to determine K$_a$ values for GroEL-ATP$_n$ reactions** | **S6** |
| **Figure S6** | **Representative spectra of GroEL-ATP$_n$ binding** | **S7** |
| **Figure S7** | **Van't Hoff plots for GroEL-ATP$_n$ binding** | **S8** |
| **Figure S8** | **Representative spectra of GroEL-ADP$_n$ binding** | **S9** |
| **Figure S9** | **Van't Hoff plots for GroEL-ADP$_n$ binding** | **S10** |

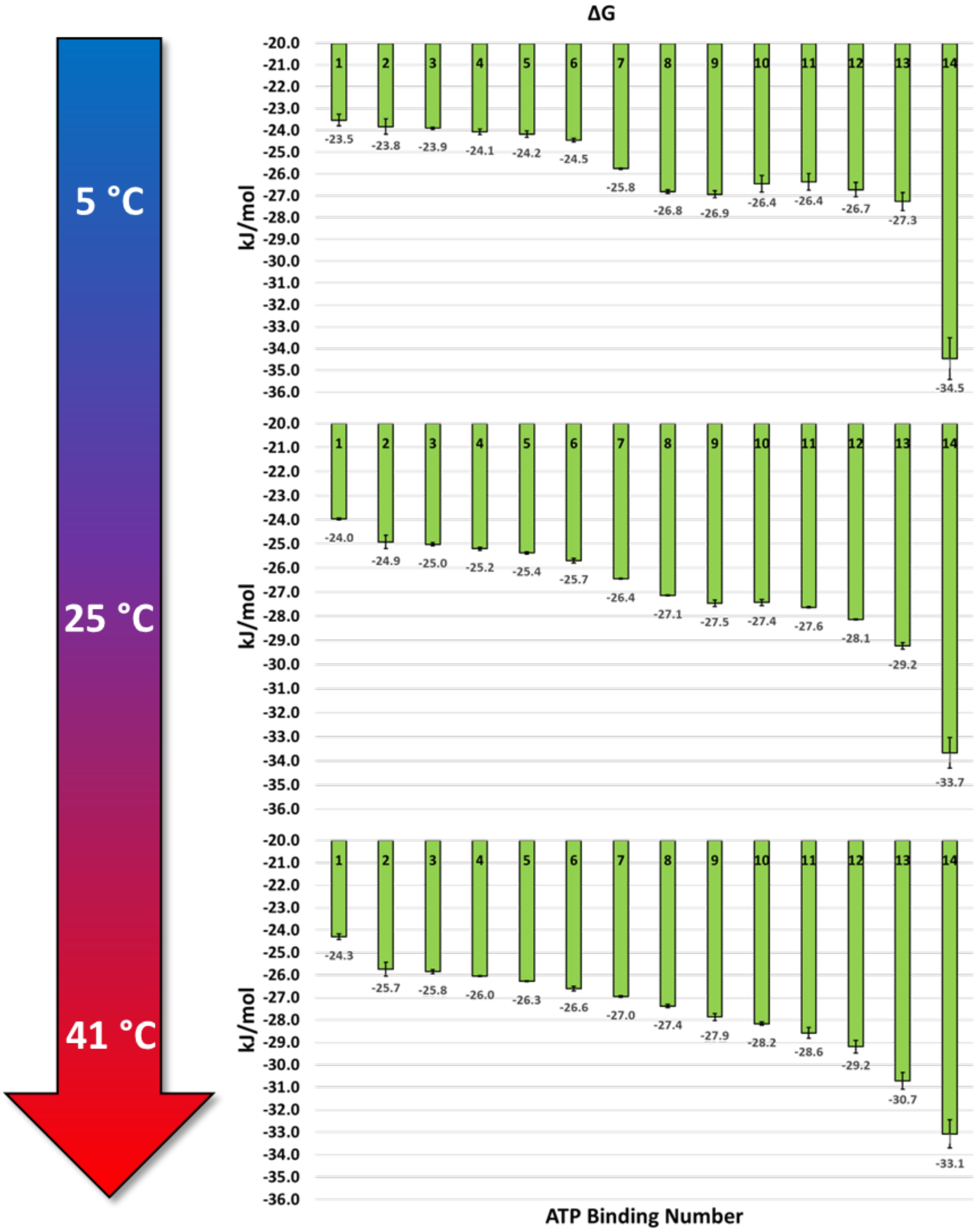


**Figure S1**. **Change in ΔG in response to change in temperature for GroEL-ATPn in EDDA.** Stacked plots displaying the changes in ΔG at various temperatures. As the solution temperature increases, the bimodality observed for the sequential ATP binding becomes diminished.

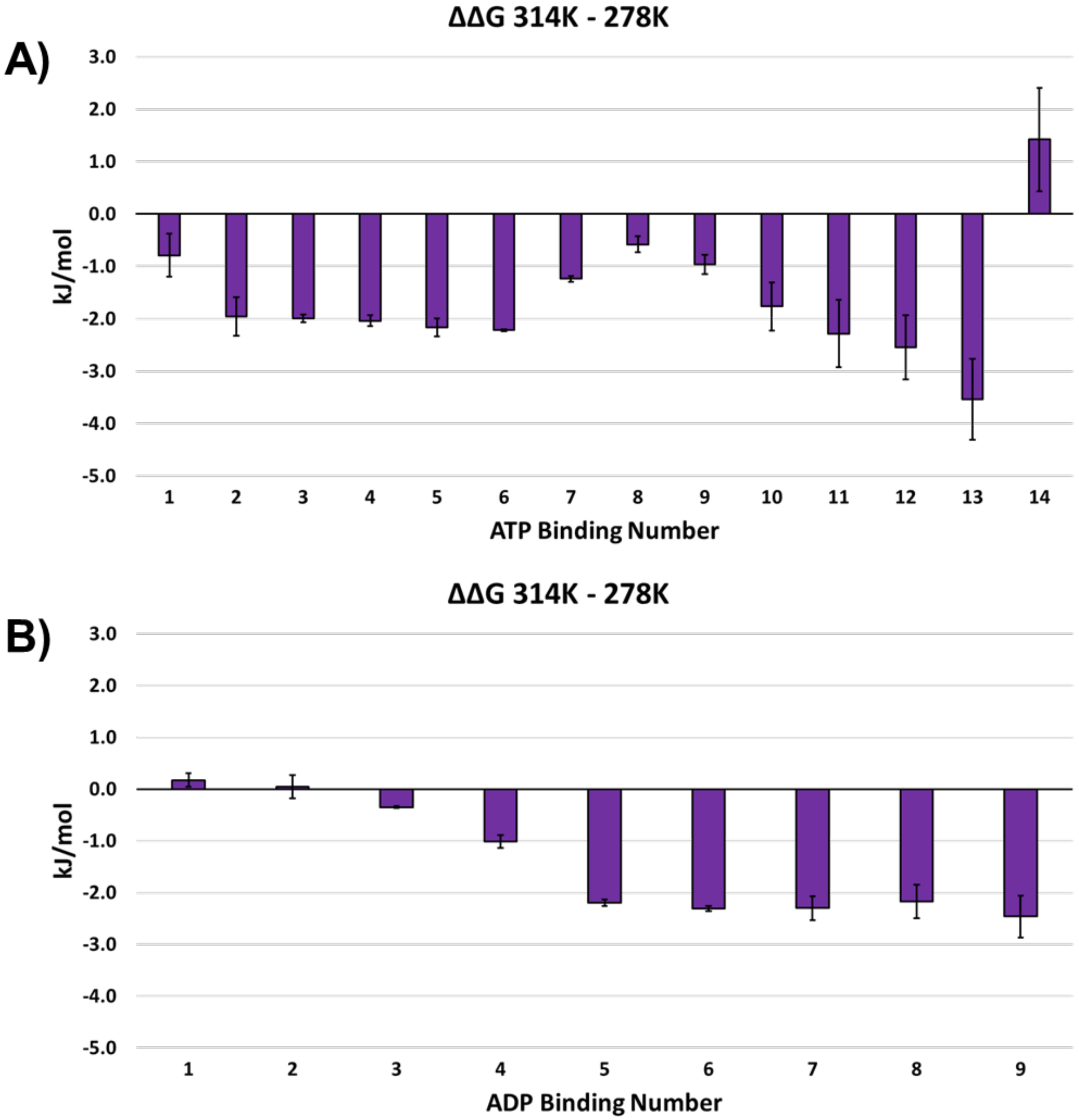


**Figure S2. ΔΔG for GroEL-ATPn in EDDA and GroEL-ADPn in AmAc.** This bar chart shows the difference in ΔG from 5 °C to 41 °C. **A)** The relative increase in the early binding reactions may explain why binding of ATP becomes less favored at higher temperatures as *cis* ring binding becomes more favored compared to *trans* ring binding. **B)** ADP binding ΔΔG values show that at higher temperatures initial ADP binding reactions become disfavored.

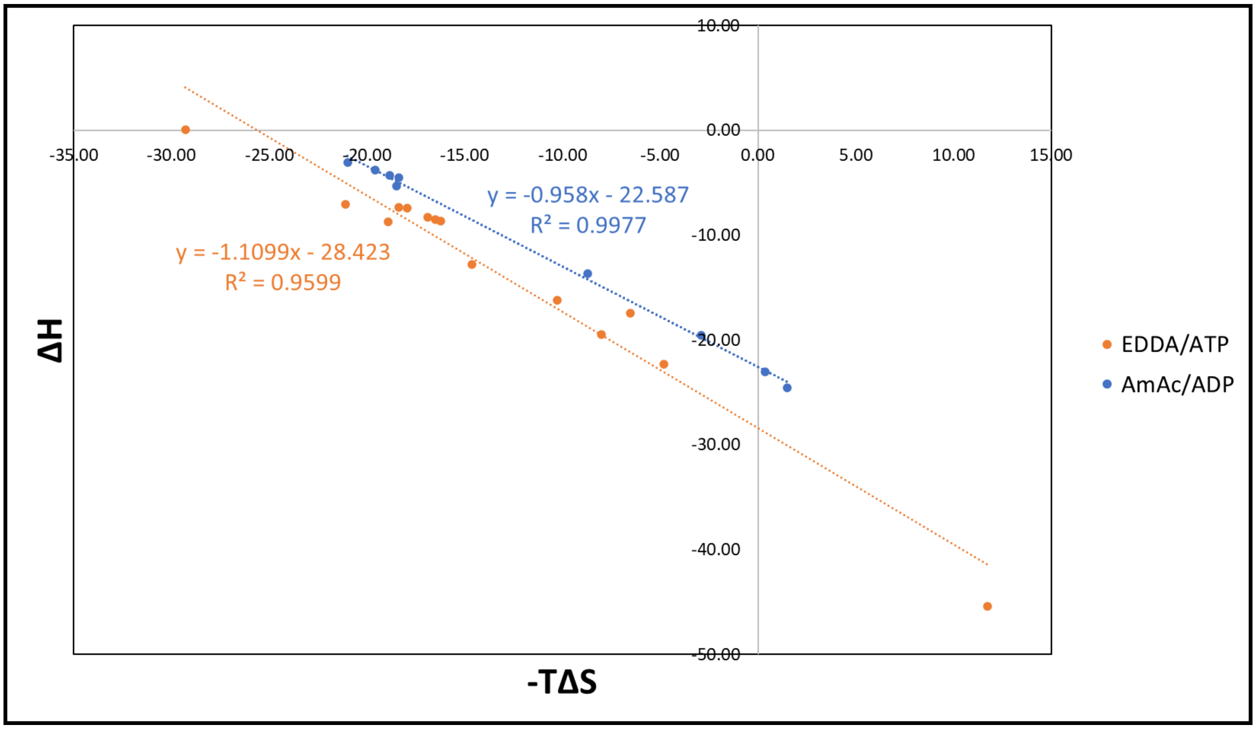


**Figure S3. Comparison of EEC for ADP and ATP binding reactions.** This plot is intended to show the EEC of entropy and enthalpy for GroEL-ATP binding in EDDA and GroEL-ADP binding in AmAc. The slope of the fit line being close to unity is a measure of the level of EEC where perfect EEC would be m = 1 (where m is the slope of the line).

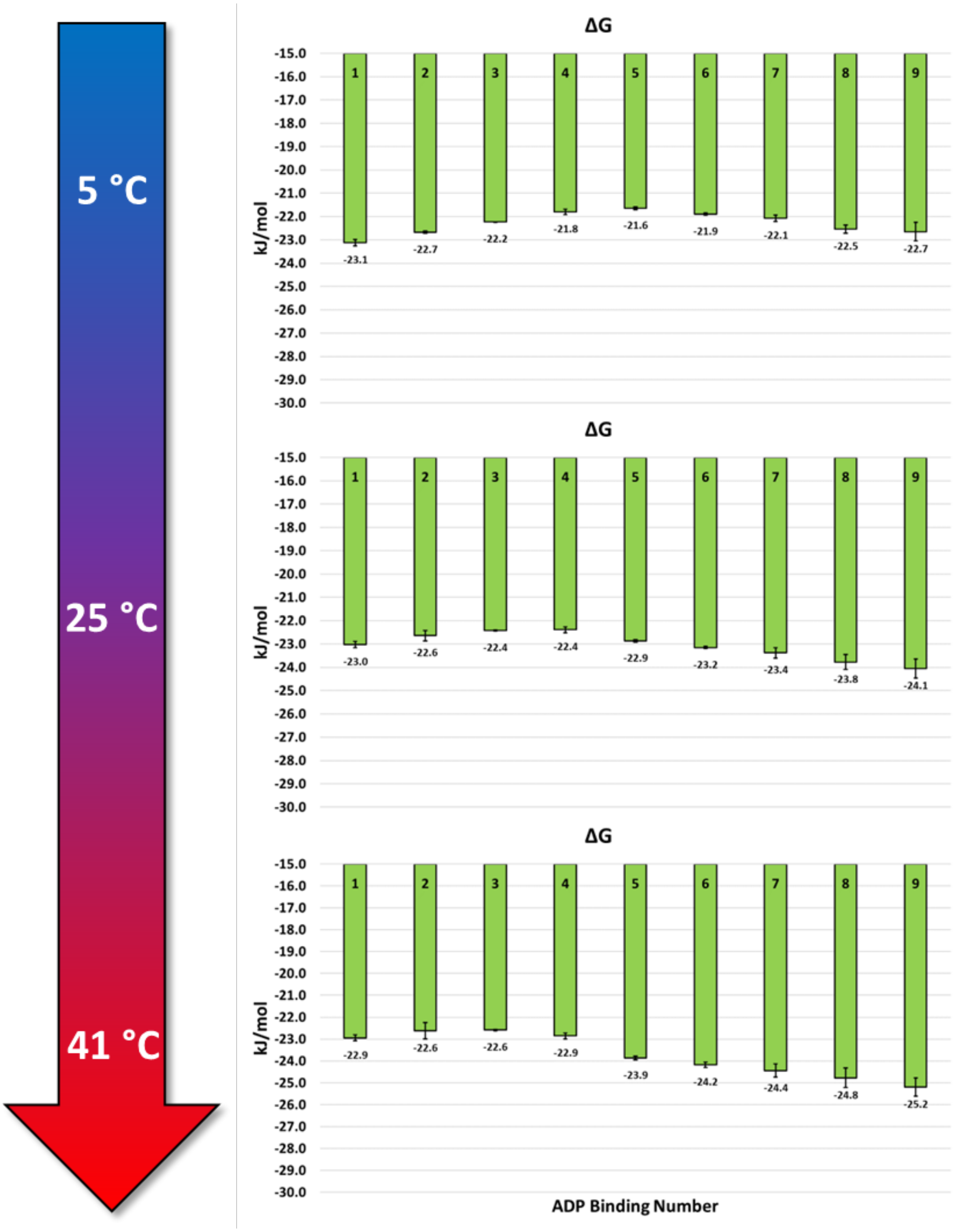


**Figure S4. Change in ΔG in response to change in temperature for GroEL-ADPn in AmAc.** Stacked plots displaying the changes in ΔG at various temperatures for the binding of ADP to GroEL.

**Table S1. ATP assignment statistics.** Average mass shift and error associated with the assignment of each [ATP + $Mg^{2+}$] (theoretical mass is ~531 Da) binding.

| ATP # | Average Mass Shift (Da) | Standard Deviation (±Da) | % Error |
|---|---|---|---|
| 1 | 548.6 | 18.4 | 3.31% |
| 2 | 551.5 | 20.0 | 3.87% |
| 3 | 536.5 | 3.9 | 1.04% |
| 4 | 538.7 | 2.4 | 1.44% |
| 5 | 539.7 | 16.0 | 1.63% |
| 6 | 522.5 | 26.6 | 1.61% |
| 7 | 531.4 | 6.6 | 0.07% |
| 8 | 539.9 | 14.5 | 1.67% |
| 9 | 542.4 | 3.0 | 2.15% |
| 10 | 539.1 | 8.0 | 1.53% |
| 11 | 519.1 | 26.4 | 2.25% |
| 12 | 523.7 | 8.3 | 1.37% |
| 13 | 527.4 | 6.9 | 0.67% |
| 14 | 551.6 | 17.6 | 3.89% |

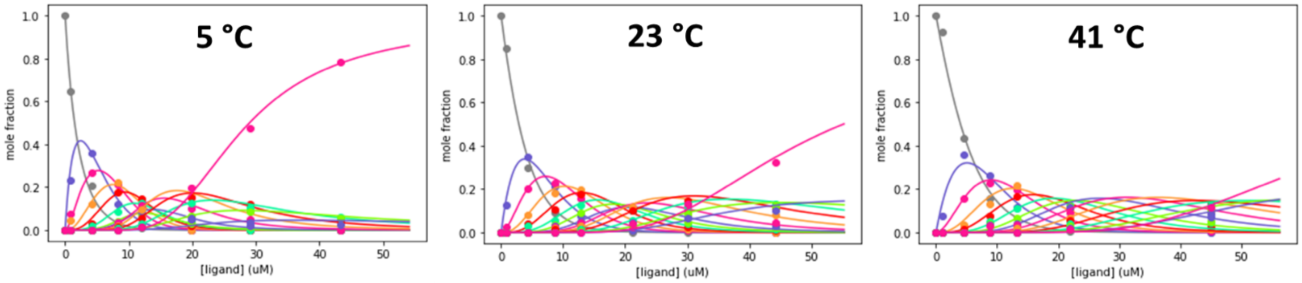


**Figure S5. Example plots used to determine Ka values for GroEL-ATPn reactions.** These mole fraction vs ATP concentration plots are used to calculate the $K_a$ values. Three temperatures are shown and the n=14 binding event (pink line) is shown to decrease in relative abundance as temperature is increased.

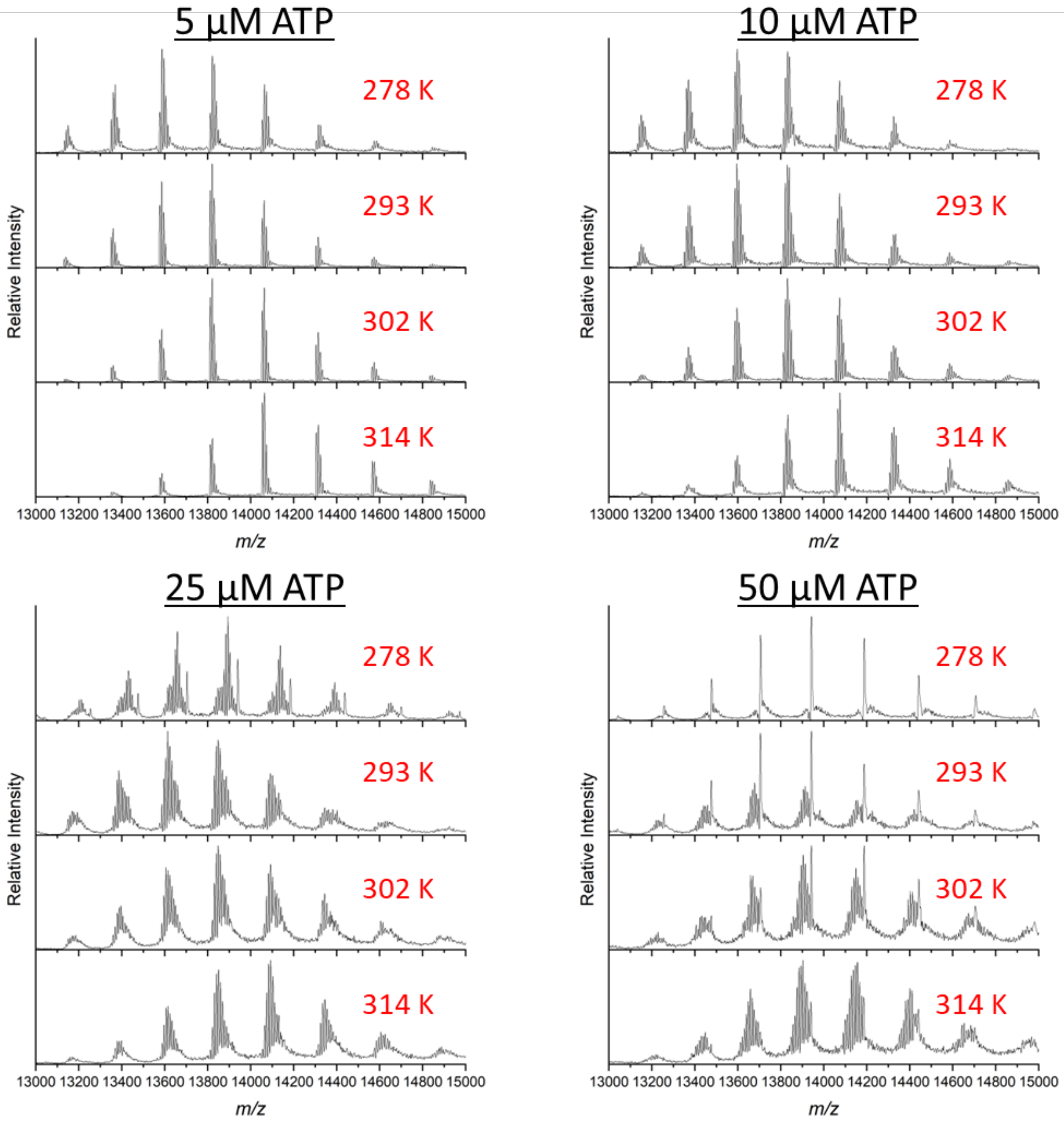


**Figure S6. Representative spectra of GroEL-ATPn binding** at various temperatures and ATP concentrations. Overall, ATP binding signals were well resolved as demonstrated above. Solution conditions are 500 nM GroEL, 200 mM EDDA, 1 mM $MgAc_2$, and various concentrations of ATP.

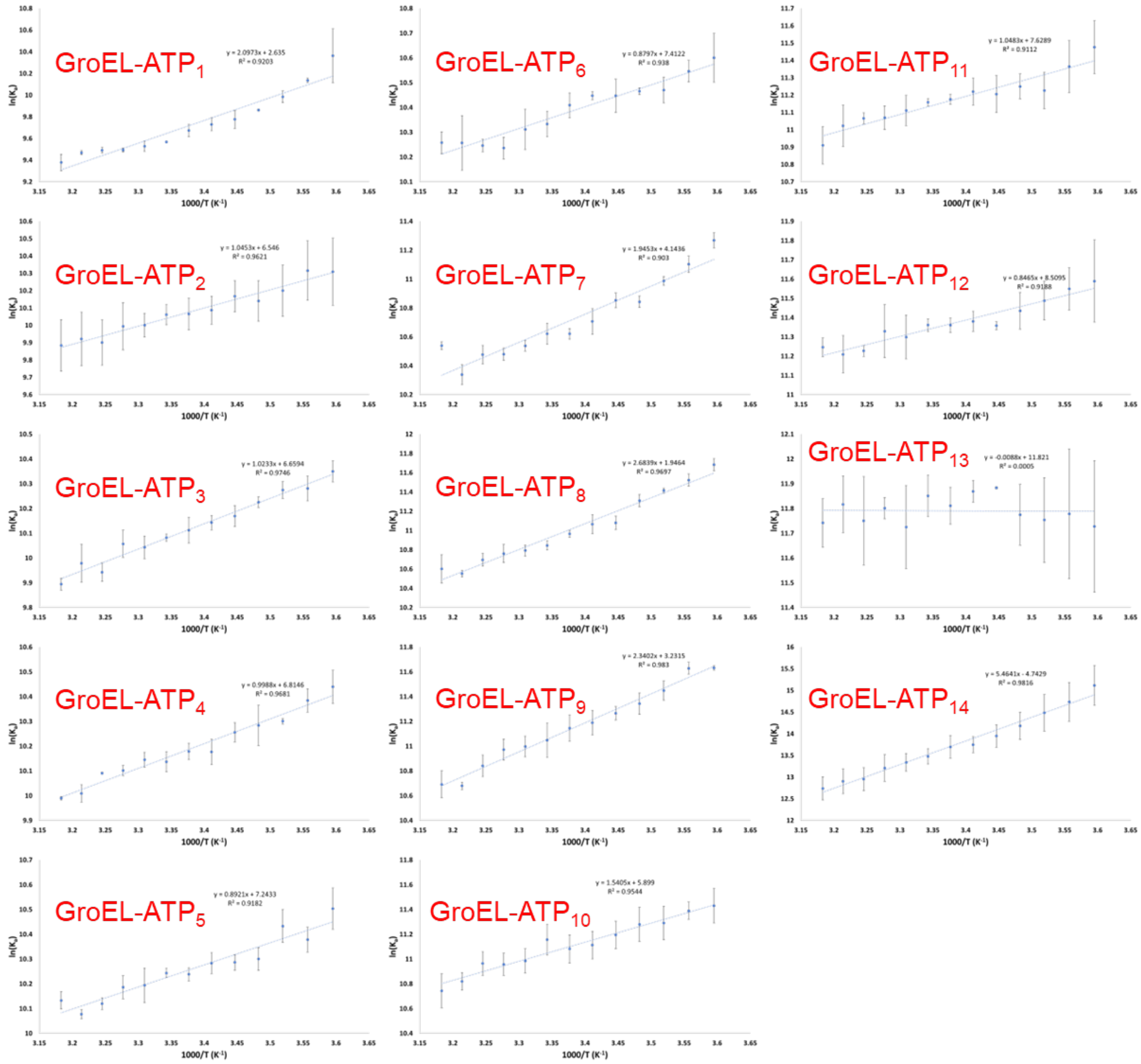


**Figure S7. Van't Hoff plots for GroEL-ATPn binding** in 200 mM EDDA ($K_a$ vs 1000/T (K)).

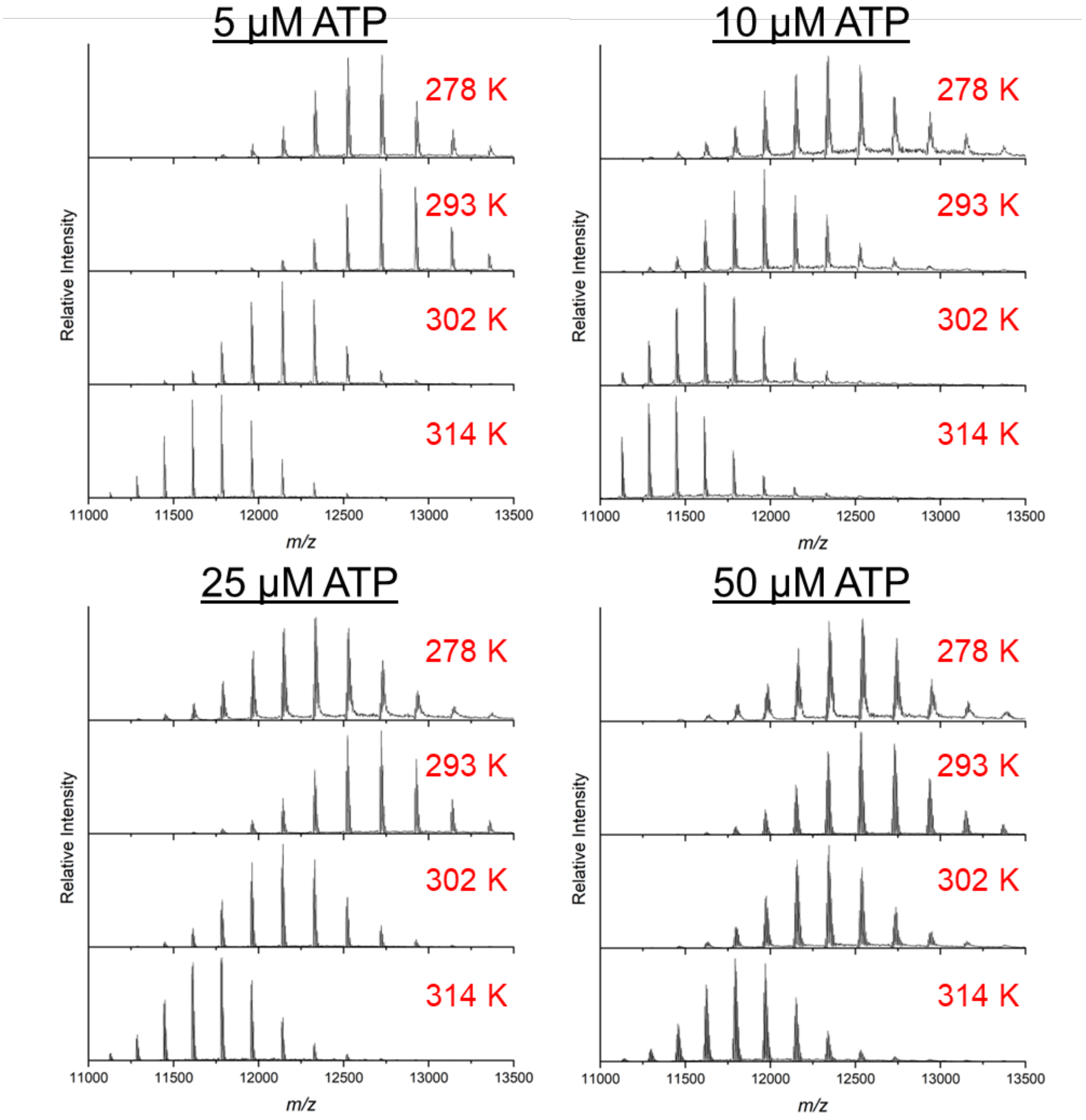


**Figure S8. Representative spectra of GroEL-ADPn binding** at various temperatures and ATP concentrations. Overall, ATP binding signals were well resolved as demonstrated above. Solution conditions are 500 nM GroEL, 200 mM AmAc, 1 mM $MgAc_2$, and various concentrations of ATP. **Note:** ATP was added to the solution but only ADP binding was observed.

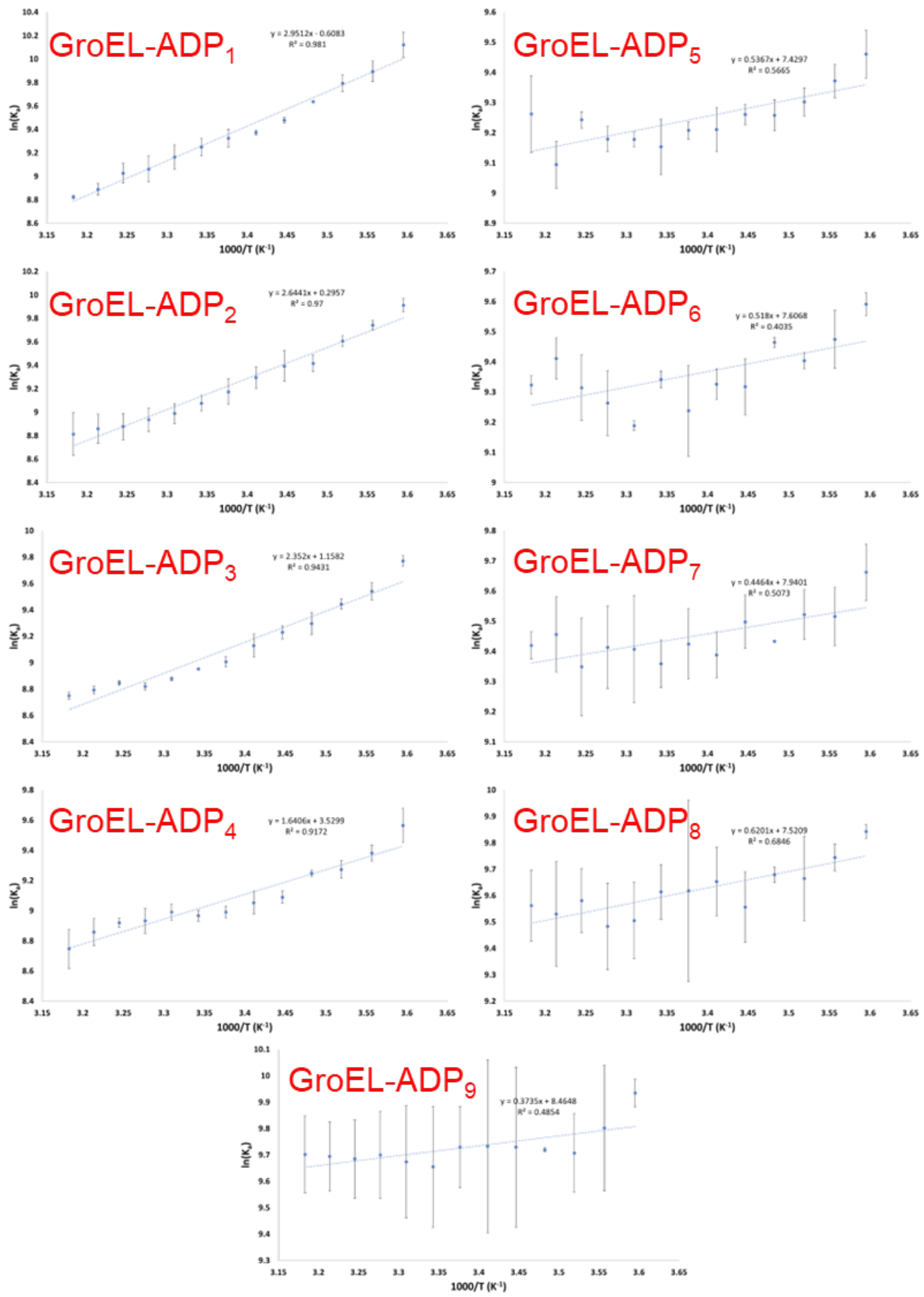


**Figure S9. Van't Hoff plots for GroEL-ADP$_n$ binding** in 200 mM AmAc ($K_a$ vs 1000/T (K)). **Note:** ATP was added to the solution but only ADP binding was observed.
